# Blast-induced mild traumatic brain injury alterations of corticotropin-releasing factor neuronal activity in the mouse hypothalamic paraventricular nucleus

**DOI:** 10.1101/2021.10.29.466475

**Authors:** Ludovic D. Langlois, Sarah Simmons, Mario Oyola, Shawn Gouty, T. John Wu, Fereshteh S. Nugent

**Affiliations:** Uniformed Services University of the Health Sciences, Department of Pharmacology and Molecular Therapeutics, Bethesda, Maryland 20814, USA; Uniformed Services University of the Health Sciences, Department of Gynecologic Surgery and Obstetrics, Bethesda, Maryland 20814, USA

**Keywords:** traumatic brain injury, blast injury, CRF, paraventricular nucleus, PVN, electrophysiology, neuronal activity, GABAergic synaptic transmission

## Abstract

Blast-induced mild traumatic brain injury (mbTBI) is the most common cause of TBI in US service members and veterans. Those exposed to TBI are at greater risk of developing neuropsychiatric disorders such as posttraumatic stress disorder, anxiety and depressive disorders, and substance use disorders following TBI [1, 2]. Previously, we have demonstrated that mbTBI increases anxiety-like behaviors in mice and dysregulates the stress at the level of corticotropin-releasing factor (CRF) neurons in the paraventricular nucleus (PVN). To expand on how mTBI may dysregulate the stress axis centrally, here PVN CRF neuronal activity was evaluated using whole cell-patch clamp recordings in hypothalamic slices from sham and mbTBI adult male CRF:tdTomato mice 7 days post-injury. We found that mbTBI generally did not affect the neuronal excitability and intrinsic membrane properties of PVN CRF neurons; this injury selectively increased the frequency of spontaneous neuronal firing of PVN CRF neurons localized to the dorsal PVN (dPVN) but not ventral PVN (vPVN). Consistently, mbTBI-induced dPVN CRF hyperactivity was associated with pre- and post-synaptic depression of spontaneous GABAergic transmission onto dPVN CRF neurons suggesting that mbTBI-induced GABAergic synaptic dysfunction may underlie dPVN CRF neuronal hyperactivity and increases in dPVN CRF signaling. The present results provide the first evidence for mbTBI-induced alterations in PVN CRF neuronal activity and GABAergic synaptic function that could mediate hypothalamic CRF dysregulation following mbTBI contributing to stress psychopathology associated with blast injury.

## 1. Introduction

Traumatic brain injury (TBI) is accounted for almost 3 million hospitalizations or admissions into the emergency room in the United States with the incidence rate increasing annually [3, 4]. Blast waves are one cause of TBI, and although they are most frequently experienced by active military personnel, civilians may also suffer from blast TBIs [5-7]. Among the non-fatal injuries, most TBIs continue to be a major source of long-lasting disabilities including impairments in cognition, mood/emotional regulation and social interactions following mTBI and contribute to the inability of affected individuals to carry out routine daily activities and to maintain important social relationships and employment [1, 8, 9]. Additionally, up to 30% of TBI patients experience neuroendocrine dysfunction [10-13] with a high incidence of stress axis [also known as the hypothalamic-pituitary-adrenal (HPA) axis] dysregulation with accompanying behavioral deficits [11, 13]. One of the major neuromodulatory stress systems that is responsive to mTBI and has significant influence on stress-related neuronal responses and affective states, is the hypothalamic PVN CRF (also known as corticotropin releasing hormone, CRH) system [14-16]. In addition to peripheral CRF endocrine signaling, recent studies suggest that central actions of CRF neurons may play an important role in regulating mood and stress modulation of behaviors. For example, it has been shown that biphasic responses of PVN CRF neuronal activity can mediate opposing behaviors of approaching appetitive stimuli or escaping from aversive stimuli [17]. Importantly, PVN CRF neurons can regulate complex behaviors in a changing environment following stress and shift innate defensive strategies [18, 19]. Previously, we have shown that blast-induced mild TBI (mbTBI)-induced neuroendocrine deficits and anxiety-like behaviors across both male and female mice include dysregulation of CRF signaling in the HPA axis but also in extrahypothalamic regions [20, 21]. Using the early response gene c-Fos immunoreactivity, as a marker for neuronal activation and retrograde Fluoro-Gold (FG) labeling of neuroendocrine-versus non-neuroendocrine-projecting CRF neurons of the PVN, we demonstrated that the central and peripheral CRF signaling is susceptible to mbTBI in both male and female mice 7 to 10 days after the blast injury. Specifically, we showed that mbTBI increased restraint stress-induced corticosterone and diminished the percentage of restraint-activated (c-Fos^+^) PVN CRF neurons in male mice but did the opposite in females. Interestingly, mbTBI exerted sex-specific differential activation of PVN neuroendocrine- (FG^+^, mostly located in ventral PVN, vPVN) and non-neuroendocrine- (FG^−^, mostly located in dPVN) projecting CRF neurons by restraint stress where the engagement of PVN non-neuroendocrine CRF neurons in mbTBI female mice was specifically lower in response to restraint stress [20]. Additionally, CRFR2 but not CRFR1 expression was affected by mbTBI in both sexes with distinct anatomical patterns of CRFR2 gene expression in stress-related PVN limbic projections [21]. Although these initial studies demonstrate that CRF stress responses from the PVN may be altered by mbTBI, it remains unclear how mbTBI dysregulates PVN CRF neuronal function in distinct anatomical and functional sub regions of the PVN. To address this, we recorded the depolarization induced neuronal excitability and spontaneous activity of CRF neurons in ventral and dorsal sub-regions of the male mouse PVN (vPVN and dPVN). We have found that mbTBI selectively increases spontaneous CRF neuronal activity within the dPVN without any significant alteration in PVN CRF neuronal excitability across dPVN and vPVN subregions. Given that the activity of distinct CRF neurons in these anatomical subregions of the PVN is shown to be biasedly modulated by some of the neurotransmitters and bioactive substances [22] and mediates endocrine versus non-neuroendocrine components of PVN CRF signaling [20, 21], our data suggest that mbTBI-induced persistent dPVN CRF dysfunction may primarily contribute to stress-related psychopathology following mbTBI.

## 2. Methods

### 2.1. Animals

7-to 9-week-old CRF:tdTomato male mice with the C57BL/6J background were generated as previously described [20] by crossing B6(Cg)-Crh^tm(cre)Zjh^/J(CRF-IRES-Cre; RRID: IMSR_JAX:012704; stock no. 012704; The Jackson Laboratory) mice and B6.Cg-Gt(ROSA) 26Sor^tm14(CAG-tdTomato)Hze^/J (Ai14; RRID: IMSR_JAX:007914; stock no. 007914; The Jackson Laboratory) mice. The mice were same-sex housed, two to three per cage, and maintained at 22°C to 25°C, 50% humidity, on a 12-hour light: 12-hour dark cycle (lights on at 0100 hours) with ad libitum access to food and water. Mice were randomly assigned to sham or mbTBI experimental group. All animal procedures were carried out in accordance with the guidelines established by the National institutes of Health (NIH) and approved by the Uniformed Services University Institutional Animal Care and Use Committee.

### 2.2. mbTBI

Mice were exposed to mbTBI under isoflurane anesthesia using the Advanced Blast Simulator (ABS; ORA Inc., Fredericksburg, VA) as previously described [20]. Sham animals were anesthetized but were not exposed to the blast injury. Immediately after sham or injury procedures, mice were observed for righting reflex and returned to the home cage for recovery. Sham and mbTBI mice were euthanized for slice preparation and electrophysiology 7 days post-injury.

### 2.3. Slice Preparation

For all electrophysiology experiments, mice were anesthetized with isoflurane, decapitated, and brains were quickly dissected and placed into ice-cold artificial cerebrospinal fluid (ACSF) containing (in mM): 126 NaCl, 21.4 NaHCO3, 2.5 KCl, 1.2 NaH2PO4, 2.4 CaCl2, 1.0 MgSO4, 11.1 glucose, and 0.4 ascorbic acid and saturated with 95% O2-5% CO2. Briefly, coronal hypothalamic slices containing PVN were cut at 250 μm and incubated in above prepared ACSF at 34°C for at least 1 h prior to electrophysiological experiments. For patch clamp recordings, slices were then transferred to a recording chamber and perfused with ascorbic-acid free ACSF at 28°C.

### 2.4. Electrophysiology

Voltage-clamp whole-cell recordings were performed from dPVN and vPVN CRF:tdTomato neurons using patch pipettes (3-6 MOhms) and a patch amplifier (MultiClamp 700B) under infrared-differential interference contrast microscopy. For all experiments, PVN CRF neurons were identified by the presence of tdTomato fluorescence which is a reliable indicator for CRF+ neurons within the PVN of CRF:tdTomato mice [23]. The anatomical boundary between dPVn and vPVN was defined by the mediolateral line through the dorsal edge of the third ventricle (3V) as previously described [22] and shown in a representative image of a hypothalamic PVN slice prepared from a CRF:tdTomato male mouse in Figure 1A. Data acquisition and analysis were carried out using DigiData 1440A, pCLAMP 10 (Molecular Devices), Clampfit, and Mini Analysis 6.0.3 (Synaptosoft, Inc.). Signals were filtered at 3 kHz and digitized at 10 kHz.

**Figure 1.**
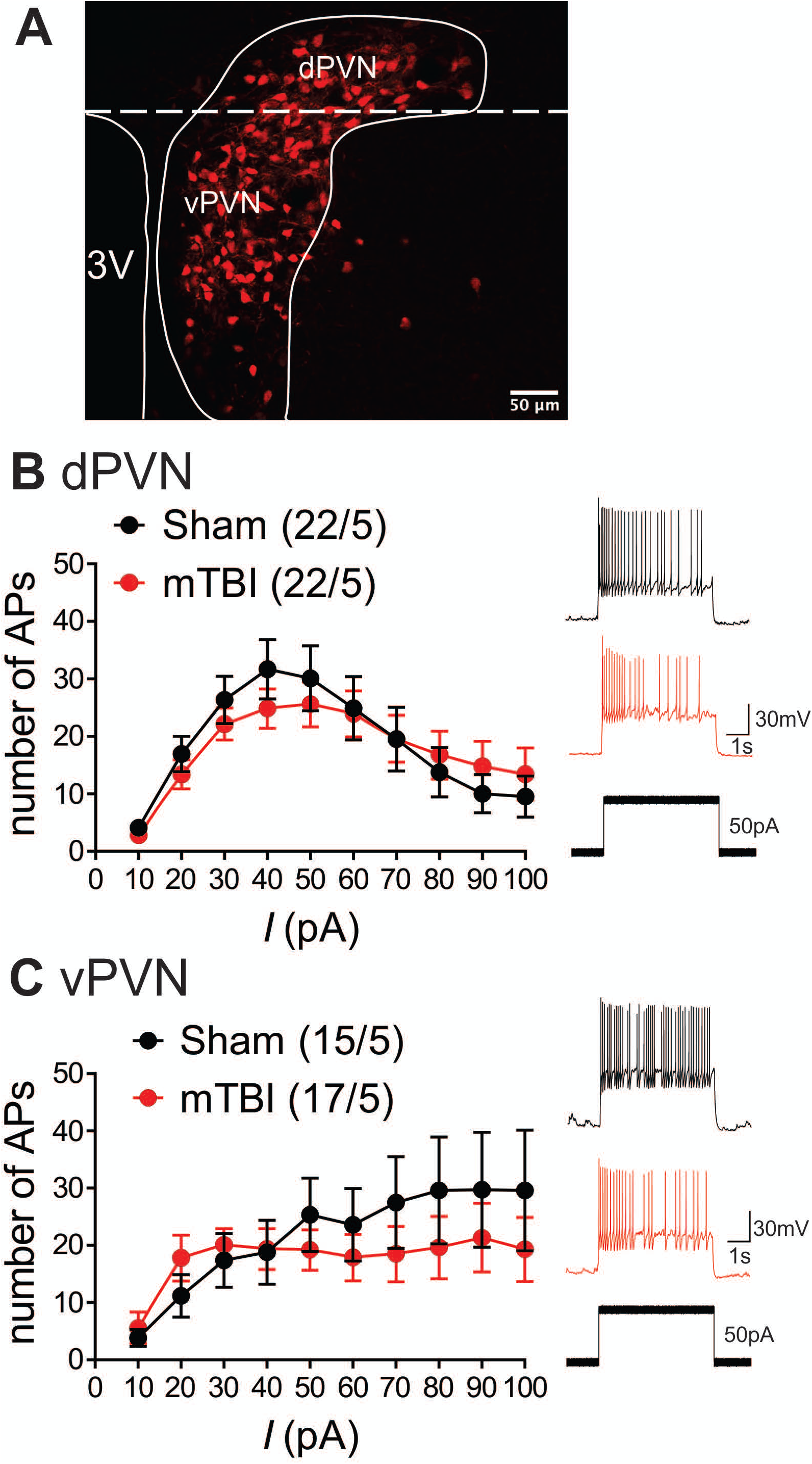
mbTBI does not alter PVN CRF neuronal excitability. **A)** Representative image showing the expression of the red fluorescent protein TdTomato in the PVN of the CRF;TdTomato mouse. Overlay is a schematic outline of the PVN labeled as the dorsal (d) PVN and ventral (v) PVN. Average number of action potentials generated across depolarizing current steps in Sham (black) and mTBI (red) mice and representative traces in response to 50pA stimulation across **B)** dPVN and **C)** vPVN CRF neurons. Group numbers presented in graph as neurons/mice.

To assess PVN CRF spontaneous neuronal activity/excitability in intact synaptic transmission, cells were patch clamped with potassium-gluconate based internal solution (130 mM K-gluconate, 15 mM KCl, 4 mM adenosine triphosphate (ATP)–Na+, 0.3 mM guanosine triphosphate (GTP)–Na+, 1 mM EGTA, and 5 mM Hepes, pH adjusted to 7.28 with KOH, osmolarity adjusted to 275 to 280 mOsm) in slices perfused with ACSF. Electrophysiological recordings of neuronal excitability, membrane properties and GABAergic transmission were performed as previously described [24, 25]. During neuronal excitability recordings in current-clamp mode, action potential (AP) generation was assessed in response to increasingly depolarizing current steps ranging from +10 to +100pA (+10 pA ea. step) while cells were kept at −67 to −70mV with manual direct current injection between pulses. Current steps were 5s in duration with 25s inter-stimulus intervals. The number of APs induced by depolarization at each intensity was counted and averaged for each experimental group at each step. Resting membrane potential (RMP) was assessed immediately after achieving whole-cell patch configuration in current clamp recordings. The hyperpolarization-activated cation current (Ih) recordings were performed in voltage-clamp in response to stepping cells from −50 mV to −100 mV (700ms duration). Input resistance (Rin) was measured at - 50pA step (5s duration) and calculated by dividing the change in voltage response by the current-pulse amplitude and presented as MΩ. AP threshold, fast after-hyperpolarizations (fAHP) and medium after-hyperpolarizations (mAHP) were assessed using clampfit and measured from the first AP at the current step that was sufficient to generate the first AP/s. Spontaneous neuronal activity and AP firing patterns were assessed in both cell-attached recordings in voltage-clamp mode at V=0mV and whole cell recording in current-clamp mode at I=0 pA for the duration of ∼1min recording. Number of APs was counted over 1 min, and spike frequency was calculated.

Whole-cell recordings of GABA_A_R-mediated spontaneous inhibitory postsynaptic currents (sIPSC) were performed in ACSF perfused with AP-V (50uM), DNQX (10 μM), and glycine receptor inhibitor (strychnine, 1 μM). Patch pipettes were filled with KCl internal solution (125 mM KCl, 2.8 mM NaCl, 2 mM MgCl2, 2mM ATP Na+, 0.3 mM GTP-Na+, 0.6 mM EGTA, and 10mM HEPES, pH adjusted to 7.28 with KOH, osmolarity adjusted to 275-280mOsm). For sIPSCs, CRF neurons were voltage-clamped at −70mV and recorded over 10 sweeps, each lasting 50s. The cell series resistance was monitored through all the experiments and if this value changed by more than 10%, data were not included.

### 2.5. Statistics

Values are presented as means ± SEM. The threshold for significance was set at *p < 0.05 for all analyses. All statistical analyses of data were performed using Graphpad, Prism 9.2. For all electrophysiological data, n represents the number of recorded cells/mouse. Mini Analysis software was used to detect and measure sIPSCs using preset detection parameters of IPSCs with an amplitude cutoff of 5 pA. Differences between sham and mbTBI mean and cumulative probabilities of sIPSC amplitude, charge transfer, tau decay and frequency were analyzed using 2-tailed unpaired Student-t-tests and Kolmogorov-Smirnov tests (KS, α = 0.05), respectively.

## 3. Results

### 3.1. mTBI does not alter PVN CRF neuronal excitability and intrinsic membrane properties independent of the anatomical locations of CRF neurons

Given our prior observations suggesting region-specific PVN CRF dysregulation, we first investigated the effects of mbTBI on PVN CRF depolarization-induced neuronal excitability and intrinsic membrane properties in intact synaptic transmission from dPVN and vPVN CRF neurons in PVN slices from sham and mbTBI male adult CRF:tdTomato mice 7 days post-injury (Figure 1 and Table 1). We found that mbTBI did not affect neuronal excitability of dPVN or vPVN CRF neurons (Figure 1B: dPVN, n=22/5 per group, 2-way ANOVA, effect of mbTBI: F (1, 420) = 0.28, P=0.59; effect of current: F (9, 420) = 8.03, P<0.0001; mbTBIxcurrent interaction: F (9, 420) = 0.47, P=0.89; Figure 2C: vPVN, n=15-17/5/group, 2-way ANOVA, effect of mbTBI: F (1, 300) = 2.11, P=0.14; effect of current: F (9, 300) = 2.32, P<0.05; mbTBIxcurrent interaction: F (9, 300) = 0.55, P=0.83). Also we found that the effects of mbTBI on intrinsic properties were negligible where mbTBI did not significantly alter RMPs, Rin, AP threshold, mAHPs although we observed a significant reduction of fAHP amplitudes only in vPVN CRF neurons (Table 1, vPVN: fAHPs, unpaired Student’s t-test, t(29)=2.37, P<0.05). Curiously, we also found that the Ih currents recorded from PVN CRF neurons in mTBI animals in general had smaller amplitudes compared to those from sham animals but only Ih currents of dPVN CRF neurons recorded from mbTBI mice showed statistically significant difference in the amplitudes in comparison to those from sham animals (Table 1, dPVN: Ih currents, unpaired Student’s t-test, t(26)=2.42, P<0.05).

**Table 1.** mbTBI had negligible impact on intrinsic membrane properties of PVN CRF neurons. Table shows the intrinsic membrane and AP properties across dPVN and vPVN CRF neurons in sham and mTBI. Unpaired Student’s t-test, *p < 0.05 noted for the statistical significance in comparisons between sham and mbTBI mice.

|  | dPVN |  | vPVN |  |
| --- | --- | --- | --- | --- |
| Property | sham | mTBI | sham | mTBI |
| RMP | -40.9 ± 1.5, n21 | -40.8 ± 1.4, n22 | -42.8 ± 1.1, n16 | -41.7 ± 1.3, n16 |
| Rin | 780.6 ± 97.3, n20 | 834.3 ± 54.45, n21 | 781.7 ± 85.49, n15 | 793.3 ± 72.22, n16 |
| AP Threshold | -31.1 ± 0.6, n18 | -32.7 ± 0.8, n21 | -34.4 ± 1.1, n14 | -32.6 ± 0.8, n16 |
| fAHP | -18.2 ± 1.2, n18 | -17.3 ± 1.0, n21 | -17.5 ± 0.9, n15 | -14.1 ± 1.2, n16<br>*P<0.05 |
| mAHP | -43.2 ± 1.5, n18 | -44.5 ± 1.5, n21 | -40.0 ± 1.5, n15 | -41.9 ± 1.8, n16 |
| Ih | -20.3 ± 5.3, n13 | -7.9 ± 1.2, n15<br>*P<0.05 | -17.2 ± 5.4, n13 | -9.1 ± 1.4, n14 |

**Figure 2.**
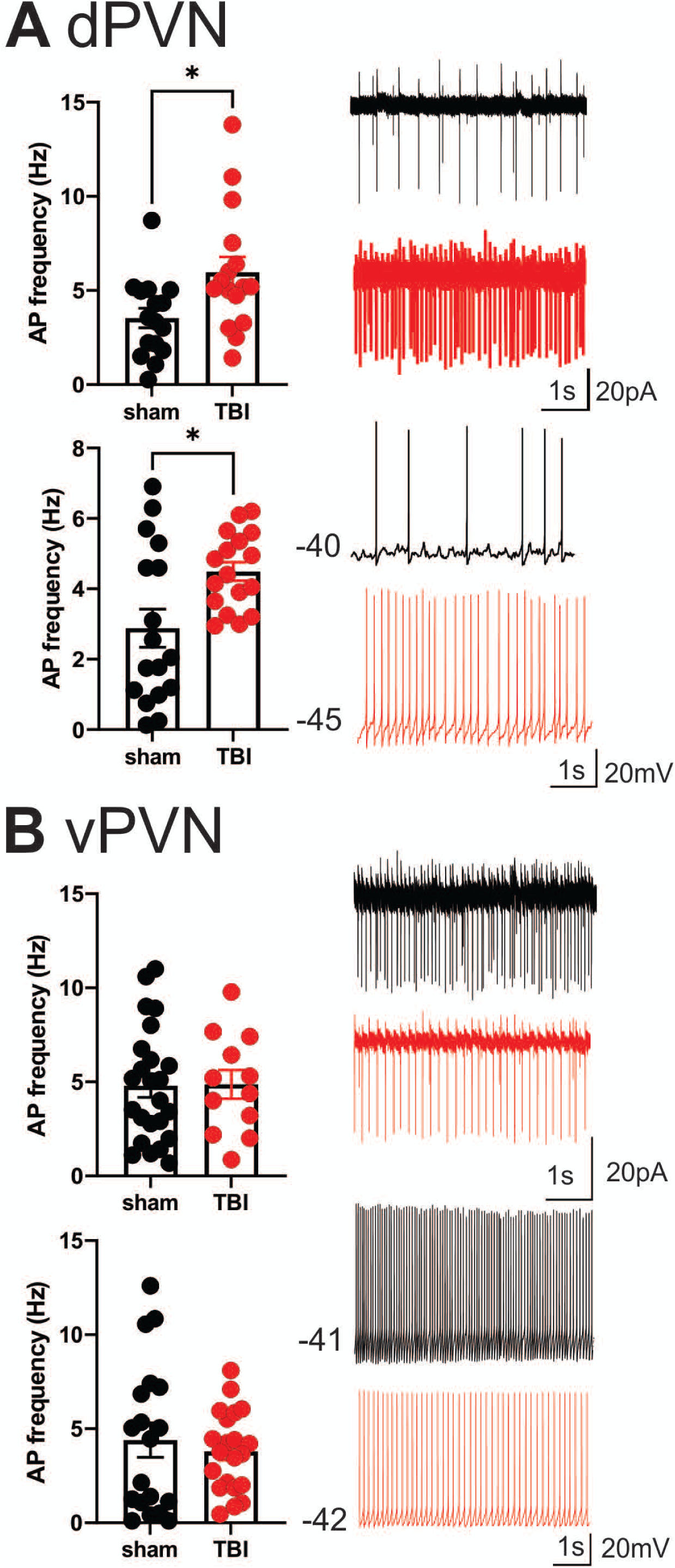
mbTBI increases spontaneous activity of dPVN CRF neurons. Representative traces, and comparison of action potential (AP) frequency under voltage-clamp (top, V = 0) cell-attached recordings and current clamp (bottom, I = 0) whole-cell recordings across sham (black) and mTBI (red) mice in **A)** dPVN and **B)** vPVN CRF neurons. n=12-24/5/Group. Unpaired Student’s t-test, *p < 0.05.

Of note, we noticed a significant phenotypic difference between dPVN versus vPVN CRF neuronal excitability of sham animals in response to depolarization where dPVN CRF neurons exhibited spike-frequency adaptation and higher AP thresholds compared to those from vPVN CRF neurons. In other words, while dPVN CRF neurons exhibited depolarization-induced blockade of AP generation in response to larger depolarizing step currents that could be due to inactivation of Na+ channels with somatic depolarization, vPVN CRF neurons with lower AP thresholds maintained high frequency firing and negligible spike frequency adaptaion (Table 1, AP threshold, unpaired Student’s t-test, t(30)=2.69, P<0.05; comparisons from sham animals in Figure 1: n=15-22/5 per group, 2-way ANOVA, effect of PVN subregion: F (1, 140) = 6.55, P<0.05; effect of current: F (9, 140) = 0.71, P=0.69; PVN subregionxcurrent interaction: F (9, 140) = 1.26, P=0.26).

### 3.2. mTBI increases dPVN but not vPVN CRF spontaneous neuronal activity

The measurements of neuronal excitability is performed in response to artificial depolarization of neurons. Therefore, to assess the spontaneous activity of PVN CRF neurons without any manipulation, we recorded cell-attached voltage clamp and whole cell current clamp recordings of spontaneous neuronal activity in dPVN and vPVN neurons in PVN slices from sham and mbTBI male adult CRF:tdTomato mice 7 days post-injury. In general, we found that CRF PVN neurons are tonically active and mbTBI did not change the number of spontaneously active neurons. However, we found that only dPVN but not vPVN CRF neurons of mbTBI mice exhibited higher mean AP firing frequency compared to those from sham mice in both voltage clamp (Figure 2A, Student’s t-test, t(30)=2.53, p<0.05) and current clamp recordings (Figure 2B, Student’s t-test, t(32)=2.67, p<0.05) suggesting that mbTBI selectively induces dPVN hyperactivity.

### 3.3. mbTBI decrease spontaneous GABAergic synaotic transmission onto dPVN CRF neurons

Given that mbTBI increase dPVN neuronal activity, we then tested the effects of mbTBI on spontaneous GABAergic neurotransmission onto dPVN neurons from sham and mbTBI male adult CRF:tdTomato mice 7 days post-injury (Figure 3). Although we did not find any significant change in the group mean sIPSC amplitude (Figure 3B), charge transfer (Figure 3C) or frequency (Figure D) following mbTBI, we observed a significant leftward shift of the CP of sIPSC amplitude (Figure 3B, KS test, p < 0.001) and charge transfer (Figure 3C, KS test, p < 0.0001) as well as a significant rightward shift of the CP of sIPSC inter-event interval (Figure 3D, KS test, p < 0.0001) following mbTBI suggesting that mbTBI-induced reduction of spontaneous GABAergic neurotransmission onto dPVN CRF neurons. Note that the group mean and CP of sIPSC Tau decay did not alter by mbTBI (Figure 3D).

**Figure 3.**
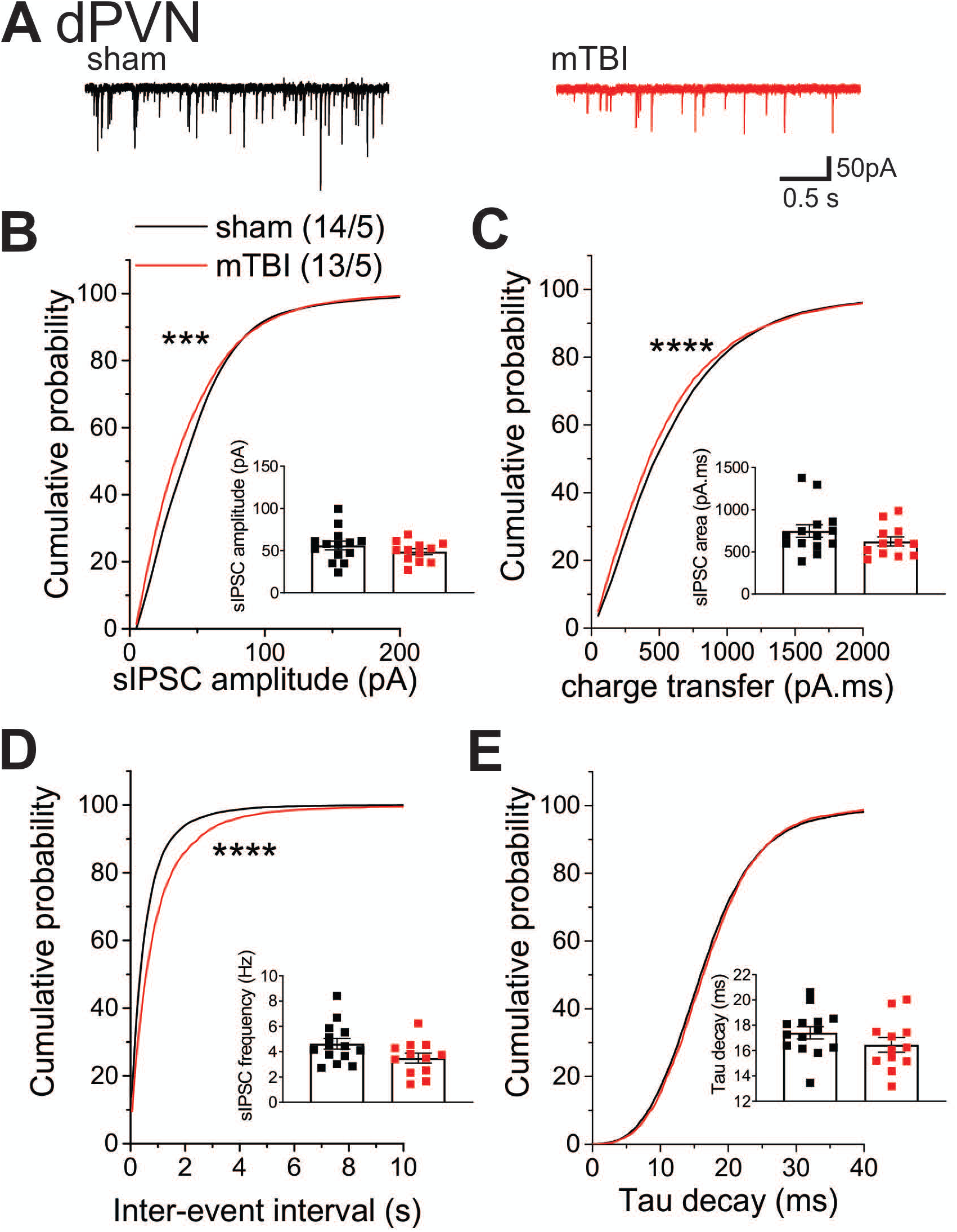
mbTBI significantly decreased spontaneous GABA_A_R-mediated synaptic transmission onto dPVN CRF neurons. **A)** Representative traces of sham (black) and mTBI (red) spontaneous IPSPs (sIPSPs). Cumulative probability (CP) curves and group mean scatter dot-plot of sIPSPs **B)** amplitude, **C)** charge transfer, **D)** inter-event interval (IEI) and **E)** Tau decay. KS test for cumulative distribution curves, *** p<0.001, **** p<0.0001. Group neuron/mice number noted in B are identical across other sIPSP graphs.

## Discussion

Previously, we showed that mbTBI may induce functional CRF dysregulation within neuroendocrine and non-neuroendocrine PVN CRF pathways [20, 21]. As a follow up to previous studies, we were interested in a functional profile of the CRF neuronal subpopulations using electrophysiology in slice preparations from sham and mbTBI male adult CRF:TdTomato mice. We found that although mbTBI did not affect the overall neuronal excitability of dPVN and vPVN CRF neurons and its effects on intrinsic membrane properties of PVN CRF neurons were negligible, it selectively resulted in an increase in spontaneous firing frequency of dPVN CRF neurons. dPVN CRF hyperactivity was also associated with decreased spontaneous GABAergic neurotransmission suggesting that mbTBI-induced GABAergic dysfunction could underlie the increased dPVN CRF signaling.

The parvocellular PVN CRF neurons mediate the physiological and behavioral stress responses of the HPA axis by releasing CRF into the median eminence which acts on the pituitary to release adrenocorticotropic hormone (ACTH). ACTH subsequently activates the synthesis and secretion of glucocorticoids, such as corticosterone from the adrenal glands to mediate stress responses. In addition to neuroendocrine PVN CRF neurons mostly located in the vPVN, the non-neuroendocrine, preautonomic dPVN CRF neurons project to the brainstem and spinal cord, and control sympathetic activity [22, 26] and may be involved in mood regulation and behavioral flexibility in response to stressors, rewarding and aversive stimuli [17-19]. Therefore, our results suggest that mbTBI-induced dysregulation of non-neuroendocrine CRF pathways from the dPVN may primarily promote anxiety-like behaviors following mbTBI. We also found that mbTBI decreased spontaneous GABAergic depression onto dPVN CRF neurons pre- and post-synaptically which could support dPVN CRF hyperactivity following mbTBI. Interestingly, a subpopulation of PVN parvocellular neurons in rats engage in a CRF feedback loop. These neurons express CRFR1 as well as CRF responsive HCN channels. CRF potentiates HCN channel activity and Ih currents and increase the neuronal activity of these PVN neurons [27]. A later study used CRF- and CRFR1-cre transgenic mouse lines with optogenetcis circuit mapping and monosynaptic tracing provided compelling evidence for the existence of intra-PVN local CRF signaling where PVN CRF neurons form synaptic connections onto preautonomic CRFR1-expressing neurons within the PVN [26]. The almost exclusive GABAergic nature of this subset of PVN CRFR1 neurons within the dPVN are rarely found to be CRF positive and allows for a local PVN inhibitory feedback. Therefore, these GABAergic CRFR1 neurons form a microcircuit within the dPVN and may respond to local release of CRF from PVN CRF neurons and in turn inhibit the activity of PVN CRF neurons. Interestingly, the ablation of PVN CRFR1 neurons also causes HPA axis hyperactivity [26] suggesting that this subset of PVN neurons provide an important inhibitory synaptic braking mechanism to prevent PVN CRF hyperexcitability. Therefore, it is possible that mbTBI may remove the intrinsic inhibitory GABAergic feedback from PVN CRFR1 neurons onto dPVN CRF neurons. Specifically, mbTBI-induced decreases in presynaptic GABA release that is evident from our sIPSC recordings in dPVN CRF neurons may indicate that mbTBI results in a loss of these intrinsic GABAergic neurons or decreases their neuronal firing although we cannot exclude the possible alteration of extrinsic GABAergic synaptic inputs to PVN CRF neurons such as the median preoptic nucleus and the posterior bed nucleus of the stria terminalis by mbTBI [28]. PVN CRF neurons receive both glutamatergic and GABAergic synaptic inputs although the proportion of fast GABAergic synaptic transmission onto PVN CRF neurons is substantially higher than other brain regions suggesting that GABAergic inhibition mediated by GABA_A_Rs and plasticity at GABAergic synapses onto PVN CRF neurons plays an important role in PVN CRF neuronal activity and restraining baseline HPA axis CRF signaling [28]. Our finding of mbTBI-induced suppression of postsynaptic GABA_A_R function in dPVN CRF neurons reflected by decreased amplitude and charge transfer of sIPSCs also suggest that mbTBI may alter or induce postsynaptic GABAergic plasticity in dPVN CRF neurons that could contribute to neuronal dysfunction although mbTBI-induced glutamatergic synaptic dysfunction in PVN CRF neurons is also likely to occur. Interestingly, we also found that mbTBI decreases in Ih currents in dPVN CRF neurons which may be a homeostatic response to dPVN CRF hyperactivity following mbTBI or an indication of intrinsic plasticity. Overall, our data suggest that mbTBI may have differential effects on distinct neuroanatomical CRF pathways where it induces persistent GABAergic synaptic dysfunction in dPVN CRF neurons to promote dPVN CRF hyperactivity thereby selectively increasing hypothalamic as well as extrahypothalamic CRF signaling and promoting anxiety-like behaviors. Given the emerging role of extrahypothalamic PVN CRF signaling, our future studies will also consider central PVN CRF projections to stress-related brain circuits that may contribute to stress psychopathology associated with this model of mTBI.

## Author Contribution

FN and TJW were responsible for the study concept and design. LL, SS, MO and SG contributed to the acquisition of animal data. FN, LL, and SS assisted with data analysis and interpretation of findings. FN, LL, SS and TJW wrote the manuscript. All authors critically reviewed content and approved final version of manuscript for submission.

## Competing interest statement

The authors have no competing interests to declare.

## Acknowledgements

The opinions and assertions contained herein are the private opinions of the authors and are not to be construed as official or reflecting the views of the Uniformed Services University of the Health Sciences or the Department of Defense or the Government of the United States. This work was supported by the National Institute of Neurological Disorders and Stroke (NIH/NINDS) Grant#R21 NS120628 to FN, and Center for Neuroscience and Regenerative Medicine and Office of Naval Research to TJW. The funding agency did not contribute to writing this article or deciding to submit it. The authors also acknowledge the USUHS Pre-Clinical Modeling and Behavior Core for supporting the studies.

## Notes

### Competing Interest Statement

The authors have declared no competing interest.

## References

1. Greer, N., et al., Prevalence and Severity of Psychiatric Disorders and Suicidal Behavior in Service Members and Veterans With and Without Traumatic Brain Injury: Systematic Review. J Head Trauma Rehabil, 2020. 35(1): p. 1–13.

2. McKee, A.C. and M.E. Robinson, Military-related traumatic brain injury and neurodegeneration. Alzheimers Dement, 2014. 10(3 Suppl): p. S242–53.

3. Taylor, C.A., et al., Traumatic Brain Injury-Related Emergency Department Visits, Hospitalizations, and Deaths - United States, 2007 and 2013. MMWR Surveill Summ, 2017. 66(9): p. 1–16.

4. Global, regional, and national burden of traumatic brain injury and spinal cord injury, 1990-2016: a systematic analysis for the Global Burden of Disease Study 2016. Lancet Neurol, 2019. 18(1): p. 56–87.

5. Hicks, R.R., et al., Neurological effects of blast injury. J Trauma, 2010. 68(5): p. 1257–63.

6. Bowen, L.N., D.F. Moore, and M.S. Okun, Is Blast Injury a Modern Phenomenon?: Early Historical Descriptions of Mining and Volcanic Traumatic Brain Injury With Relevance to Modern Terrorist Attacks and Military Warfare. Neurologist, 2016. 21(2): p. 19–22.

7. Helmick, K.M., et al., Traumatic brain injury in the US military: epidemiology and key clinical and research programs. Brain Imaging Behav, 2015. 9(3): p. 358–66.

8. Silver, J.M., T.W. McAllister, and D.B. Arciniegas, Depression and cognitive complaints following mild traumatic brain injury. Am J Psychiatry, 2009. 166(6): p. 653–61.

9. Wong, E.C., et al., Mental health treatment experiences of U.S. service members previously deployed to Iraq and Afghanistan. Psychiatr Serv, 2013. 64(3): p. 277–9.

10. Molaie, A.M. and J. Maguire, Neuroendocrine Abnormalities Following Traumatic Brain Injury: An Important Contributor to Neuropsychiatric Sequelae. Front Endocrinol (Lausanne), 2018. 9: p. 176.

11. Krahulik, D., et al., Dysfunction of hypothalamic-hypophysial axis after traumatic brain injury in adults. J Neurosurg, 2010. 113(3): p. 581–4.

12. Lieberman, S.A., et al., Prevalence of neuroendocrine dysfunction in patients recovering from traumatic brain injury. J Clin Endocrinol Metab, 2001. 86(6): p. 2752–6.

13. Hoffman, A.N. and A.N. Taylor, Stress reactivity after traumatic brain injury: implications for comorbid post-traumatic stress disorder. Behav Pharmacol, 2019. 30(2 and 3-Spec Issue): p. 115–121.

14. Kosari-Nasab, M., et al., The blockade of corticotropin-releasing factor 1 receptor attenuates anxiety-related symptoms and hypothalamus-pituitary-adrenal axis reactivity in mice with mild traumatic brain injury. Behav Pharmacol, 2019. 30(2 and 3-Spec Issue): p. 220-228.

15. Fox, L.C., et al., Differential effects of glucocorticoid and mineralocorticoid antagonism on anxiety behavior in mild traumatic brain injury. Behav Brain Res, 2016. 312: p. 362–5.

16. McCorkle, T.A., J.R. Barson, and R. Raghupathi, A Role for the Amygdala in Impairments of Affective Behaviors Following Mild Traumatic Brain Injury. Front Behav Neurosci, 2021. 15: p. 601275.

17. Kim, J., et al., Rapid, biphasic CRF neuronal responses encode positive and negative valence. Nature Neuroscience, 2019. 22(4): p. 576–585.

18. Füzesi, T., et al., Hypothalamic CRH neurons orchestrate complex behaviours after stress. Nature Communications, 2016. 7(1): p. 11937.

19. Daviu, N., et al., Paraventricular nucleus CRH neurons encode stress controllability and regulate defensive behavior selection. Nature Neuroscience, 2020. 23(3): p. 398–410.

20. Russell, A.L., et al., Differential Responses of the HPA Axis to Mild Blast Traumatic Brain Injury in Male and Female Mice. Endocrinology, 2018. 159(6): p. 2363–2375.

21. Russell, A.L., R.J. Handa, and T.J. Wu, Sex-Dependent Effects of Mild Blast-induced Traumatic Brain Injury on Corticotropin-releasing Factor Receptor Gene Expression: Potential Link to Anxiety-like Behaviors. Neuroscience, 2018. 392: p. 1–12.

22. Mukai, Y., et al., Identification of substances which regulate activity of corticotropin-releasing factor-producing neurons in the paraventricular nucleus of the hypothalamus. Scientific Reports, 2020. 10(1): p. 13639.

23. Wamsteeker Cusulin, J.I., et al., Characterization of corticotropin-releasing hormone neurons in the paraventricular nucleus of the hypothalamus of Crh-IRES-Cre mutant mice. PLoS One, 2013. 8(5): p. e64943.

24. Authement, M.E., et al., A role for corticotropin-releasing factor signaling in the lateral habenula and its modulation by early-life stress. Sci Signal, 2018. 11(520).

25. Simmons, S.C., et al., Early life stress dysregulates kappa opioid receptor signaling within the lateral habenula. Neurobiol Stress, 2020. 13: p. 100267.

26. Jiang, Z., S. Rajamanickam, and N.J. Justice, Local Corticotropin-Releasing Factor Signaling in the Hypothalamic Paraventricular Nucleus. J Neurosci, 2018. 38(8): p. 1874–1890.

27. Qiu, D.L., et al., Corticotrophin-releasing factor augments the I(H) in rat hypothalamic paraventricular nucleus parvocellular neurons in vitro. J Neurophysiol, 2005. 94(1): p. 226–34.

28. Bains, J.S., J.I. Cusulin, and W. Inoue, Stress-related synaptic plasticity in the hypothalamus. Nat Rev Neurosci, 2015. 16(7): p. 377–88.

